# What do I do now? Intolerance of uncertainty is associated with discrete patterns of anticipatory physiological responding to different uncertain contexts

**DOI:** 10.1101/527317

**Authors:** Jayne Morriss

## Abstract

Heightened physiological responses to uncertainty are a common hallmark of anxiety disorders. Many separate studies have examined the relationship between individual differences in intolerance of uncertainty (IU) and physiological responses to uncertainty during different contexts. Despite this, little is known about the extent to which physiological responses during different uncertain contexts covary within individuals based on IU. Anticipatory physiological responses to uncertainty were assessed in three different contexts (associative threat learning, basic threat uncertainty, decision-making) within the same sample (n = 45). During these tasks, behavioural responses (i.e. reaction times, choices), skin conductance and corrugator supercilli activity were recorded. In addition, self-reported IU and trait anxiety were measured. IU made different contributions to the physiological measures during each task. IU was found to modulate both skin conductance and corrugator supercilii activity for the associative threat learning and decision-making contexts. However, trait anxiety was found to modulate corrugator supercilii activity during the basic threat uncertainty context. Ultimately, this research helps us further tease apart the role of IU in different uncertain contexts, which will be relevant for future IU-related models of psychopathology.

## 1. Introduction

On a daily basis we are confronted with uncertainty about the outcome of future events. In some uncertain situations we may be able to predict the outcome, or weigh up the options and make a choice, whilst in other uncertain situations we have no control of the outcome, whatever it may be. Individuals who are high in intolerance of uncertainty (IU) are described as having a ‘dispositional incapacity to endure the aversive response triggered by the perceived absence of salient, key, or sufficient information, and sustained by the associated perception of uncertainty’ (Carleton, 2016b, p 31). IU has been linked to many mental health disorders that have an anxiety component e.g. anxiety, depression and obsessive compulsive disorder (Gentes & Ruscio, 2011; McEvoy & Mahoney, 2012). On this basis the study of IU has gained substantial momentum in recent years and now sits at the forefront of anxiety research (Grupe & Nitschke, 2013; Tanovic, Gee, & Joormann, 2018). Initial work shows that individuals high in IU, relative to individuals low in IU, display heightened physiological and neural activity to uncertainty across several different domains i.e. threat, reward, and error monitoring (for review see Tanovic, Gee & Joorman, 2018). The most popular areas of research for examining the neural correlates of uncertainty and IU have been in the following contexts: (1) associative threat learning, (2) basic threat uncertainty, and (3) decision-making (for review see, Morriss, Gell & van Reekum, 2018).

There is robust evidence for the role of IU in associative learning, particularly when associations need to be updated from threat to safe, as in threat extinction (Chin, Nelson, Jackson, & Hajcak, 2016; Dunsmoor, Campese, Ceceli, LeDoux, & Phelps, 2015; Lucas, Luck, & Lipp, 2018; Morriss, Christakou, & Van Reekum, 2015, 2016; Morriss, Macdonald, & van Reekum, 2016). The context of threat extinction is inherently uncertain, as at the start of extinction the contingencies are unknown. This likely induces uncertainty-induced anxiety in individuals with high IU. For example, after 100% reinforcement, high IU, relative to low IU individuals have been found to show generalized skin conductance response and amygdala activity across threat and safety cues during early extinction, and to show continued skin conductance responding and amygdala activity to threat versus safety cues during late extinction (Morriss, Christakou, & van Reekum, 2015, 2016). Moreover, after 50% reinforcement, high IU has been found to be associated with generalized skin conductance responding to stimuli that vary in similarity to the learned threat cue during extinction (Morriss, Macdonald, & van Reekum, 2016).

The evidence for the role of IU on basic threat uncertainty is mixed (Bennett, Dickmann, & Larson, 2018; Gole, Schäfer, & Schienle, 2012; Grupe & Nitschke, 2011; Schienle, Köchel, Ebner, Reishofer, & Schäfer, 2010; Somerville et al., 2013). In the basic threat uncertainty tasks the uncertainty and valence parameters are known, thus the goal of the participant is to tolerate the potential for an uncertain aversive event. For example, participants will be presented cues that predict either negative or neutral pictures (Grupe & Nitschke, 2011), or will follow a predictable or unpredictable countdown to a negative or neutral picture (Somerville et al., 2013). Therefore, these tasks are labelled as ‘basic’ because there is no associative learning or action to be made. The need to tolerate uncertain threatening outcomes may invoke some uncertainty-induced anxiety in individuals scoring high in IU. During these tasks, individuals high in IU, relative to low IU have been shown to exhibit heightened amygdala activity to cues conveying uncertainty (Schienle et al., 2010) and to aversive pictures following unpredictable countdowns (Somerville et al., 2013). However, for these basic uncertainty tasks there is a paucity of significant results for an effect of IU on anticipatory physiological responses (Bennett et al., 2018; Grupe & Nitschke, 2011).

There is a small albeit emerging literature on the role of IU in decision-making. The decision-making tasks that have been used to examine IU vary substantially. In the majority of the decision-making tasks the uncertainty and valence parameters are known, and the goal of the participant is to make optimal decisions in order to gain reward and avoid loss. In these tasks decisions under uncertainty must be made and cannot be avoided, thus these tasks may invoke some uncertainty-induced anxiety in individuals scoring high in IU. Individuals high in IU have been found to report more distress (Jacoby, Abramowitz, Reuman, & Blakey, 2016; Jacoby, Reuman, Blakey, Hartsock, & Abramowitz, 2017) and make more draws to decision on the beads task (Jacoby, Abramowitz, Buck, & Fabricant, 2014; Ladouceur, Talbot, & Dugas, 1997). In addition, a few studies have demonstrated that high IU individuals are more likely to choose immediate smaller rewards over waiting for larger rewards (Carleton et al., 2016; Luhmann, Ishida, & Hajcak, 2011; Tanovic, Hajcak, & Joormann, 2018). Taken together these findings suggest that individuals scoring high in IU will seek more information to reduce uncertainty and will not wait to make a decision when there is no additional information about the uncertain outcome. Despite this progress, there are a lack of studies that have examined how IU may impact anticipatory physiological responses during decision-making.

What is noticeable from the areas of research outlined above is that they all contain aspects of anticipation (Morriss, Gell, & van Reekum, 2018). Whether it be the anticipation of: (1) a learned uncertain threat outcome, (2) a known uncertain threat outcome or (3) making a decision under uncertainty. Despite the advances in the field, there is a scarcity of research examining the extent to which anticipatory physiological responses and individual differences in IU are shared or discrete depending on the type of uncertain context. This makes it difficult to assess the generalizability or specificity of IU-related physiological profiles and their relevance to psychopathology models and aetiology (Shihata, McEvoy, Mullan, & Carleton, 2016). Based on this, the following study attempted to fill in some of the gaps in the literature by measuring anticipatory physiological responses to different uncertain contexts: an associative threat learning task with acquisition and extinction phases (Morriss et al., 2015; Morriss, Christakou, et al., 2016); a basic uncertainty task with cues that predict negative or neutral pictures (Gole et al., 2012; Grupe & Nitschke, 2011; Schienle et al., 2010); a decision-making task where uncertain stimuli are categorised in the absence of reward or loss.

The associative threat learning and basic uncertainty tasks were similar to those that had already been designed to assess IU. Both these tasks included anticipated aversive outcomes, as previous literature using these tasks has focused on the interaction between IU, levels of uncertainty, and threat (Grupe & Nitschke, 2013; Tanovic, Gee, et al., 2018). The decision-making task was designed to assess the anticipation of making a decision in the absence of valenced outcomes. The majority of previous studies on IU and decision-making have focused on reward and loss. However, it is unknown whether decision-making under uncertainty in the absence of valenced outcomes is aversive enough to modulate anticipatory physiological responses. Indeed, previous research has observed IU to modulate other psychological mechanisms such as attention in the absence of threat (Fergus, Bardeen, & Wu, 2013; Fergus & Carleton, 2016).

Throughout the tasks anticipatory physiological responses were measured using skin conductance and corrugator supercilli activity. Skin conductance response is thought to capture arousal (Dawson, Schell, & Filion, 2000) and corrugator supercilli activity is thought to capture valence (Tassinary, Cacioppo, & Vanman, 2007). For example: (1) greater skin conductance response is observed for both negative and positive pictures, relative to neutral pictures, and (2) greater corrugator supercilli activity is observed for negative, relative to positive and neutral pictures (Bradley & Lang, 2000). The advantage of recording both skin conductance response and corrugator supercilli activity is that it may reveal whether there are specific aspects of arousal and valence related to IU and anticipation during the different uncertain contexts.

Alongside the physiological measures, ratings of threat vs. safety cues were recorded for the associative learning task, and choices and reaction times were recorded for the decision-making task. In addition, self-reported IU and trait anxiety (STAI) (Spielberger, Gorsuch, Lushene, Vagg, & Jacobs, 1983) were measured. STAI is a popular measure for self-reported anxiety and has been used in associative learning, basic threat uncertainty and decision-making literatures (Grupe & Nitschke, 2013; Hartley & Phelps, 2012; Lonsdorf & Merz, 2017). To assess the specificity of IU during different uncertain contexts, it is useful to contrast it with other self-reported anxiety measures such as STAI (for discussion see Morriss, Christakou & van Reekum, 2016).

For the associative learning task it was predicted that all participants would display greater expectancy ratings, skin conductance and corrugator supercilli activity to learned threat (CS+) vs. safety (CS-) cues during acquisition, and that the differential response to the CS+ vs. CS-for skin conductance and corrugator supercilli activity would reduce over time during extinction. Given previous research, it was predicted that individuals scoring higher in IU would show heightened skin conductance and corrugator supercilli activity during extinction (Morriss et al., 2015; Morriss, Christakou, et al., 2016; Morriss, Macdonald, et al., 2016). No predictions were made for IU and expectancy ratings during the associative learning task, due to the lack of findings in previous studies.

For the basic threat uncertainty task it was predicted that all participants would exhibit larger skin conductance and corrugator supercilli activity to: (1) cues predicting certain negative pictures vs. cues predicting uncertain pictures and cues predicting certain neutral pictures, and (2) negative pictures following certain cues vs. negative pictures following uncertain cues and neutral pictures following uncertain/certain cues. Moreover, it was predicted that higher IU would be associated with greater skin conductance and corrugator supercilli activity to the uncertain cue and to negative pictures following an uncertain cue, relative to the other conditions.

Lastly, for the decision-making task, it was predicted that all participants would display relatively accurate choices based on probability. In addition, it was predicted that all participants would show larger reaction times, skin conductance and corrugator supercilli activity when making decisions on uncertain stimuli vs. certain stimuli. Furthermore, it was predicted that individuals scoring higher in IU relative to lower IU would display larger reaction times, skin conductance and corrugator supercilli activity when making decisions on uncertain stimuli vs. certain stimuli.

If IU is related to anticipatory physiological responses across uncertain contexts then IU likely serves to modulate anticipatory mechanisms generally. Alternatively, if IU is only related to anticipatory responses in some uncertain contexts then it suggests that IU may serve to modulate anticipatory mechanisms distinctly depending on the uncertain context i.e. associative threat learning, basic threat uncertainty or decision-making under uncertainty.

## 2. Method

### 2.1 Participants

Forty-five volunteers (M age = 23 years, SD age = 4.23 years; 31 females and 14 males; 33 Europeans, 5 Asian, 4 African/Afro Caribbean, 3 Mixed) took part in the study. The sample size was based on previous experiments that have examined IU-related differences in associative learning using psychophysiological measures (Chin, Nelson, Jackson, & Hajcak, 2016; Dunsmoor, Campese, Ceceli, LeDoux, & Phelps, 2015; Lucas, Luck, & Lipp, 2018; Morriss, Christakou, & van Reekum, 2016; Morriss, MacDonald, & van Reekum, 2016). All participants had normal or corrected to normal vision. Participants provided written informed consent and received £10 for their participation. Advertisements and word of mouth were used to recruit participants from the University of Reading and local area. No exclusion criteria were used. Two participants withdrew from the aversive learning task and one withdrew from the basic threat uncertainty task. There was a recording error for one participant on the aversive learning task. The procedure was approved by the University of Reading Research Ethics Committee.

### 2.2 Procedure

On the day of the experiment participants arrived at the laboratory and were informed on the experimental procedures. Firstly, participants were seated in the testing booth and asked to complete and sign a consent form as an agreement to take part in the study. Secondly, participants completed questionnaires on a computer (see 2.4 below). Thirdly, electromyography sensors were attached to the left corrugator supercilli and physiological sensors were attached to the participants’ non-dominant hand. The tasks (see 2.3 below) were presented in a counterbalanced order on a computer, whilst skin conductance, interbeat interval and behavioural ratings were recorded. Participants were instructed to maintain attention to the tasks and to stay as still as possible. The experiment took approximately 60 minutes in total.

### 2.3 Tasks

All tasks were designed using E-Prime 2.0 software (Psychology Software Tools Ltd, Pittsburgh, PA). Visual stimuli were presented at a 60 Hz refresh rate on an 800 × 600 pixel computer screen. Participants sat approximately 60 cm from the screen.

#### 2.3.1 Associative threat learning task

The experimental design for the associative threat learning task was identical to previous work (Morriss & van Reekum, Under Review). Participants were required to passively view cues that predicted a threat or safe outcome. Participants did not receive instructions about the threat/safe contingencies and were simply told to pay attention to the squares and sounds.

Visual stimuli were blue and yellow squares with 183 × 183 pixel dimensions. The aversive sound stimulus was presented through headphones. The sound consisted of a threat inducing female scream used in previous experiments (Morriss et al., 2015; Morriss et al., 2016). The volume of the sound was standardized across participants by using fixed volume settings on the presentation computer and was verified by an audiometer prior to each session (90 dB).

The task comprised of two learning phases: acquisition and extinction. Both acquisition and extinction consisted of two blocks each. In acquisition, one of the coloured squares (blue or yellow) was paired with the aversive 90 dB sound 50% of the time (CS+), whilst the other square (yellow or blue) was presented alone (CS-). During extinction, both the blue and yellow squares were presented in the absence of the US.

The acquisition phase consisted of 24 trials (6 CS+ paired, 6 CS+ unpaired, 12 CS-) and the extinction phase 32 trials (16 CS+ unpaired, 16 CS-). Experimental trials were pseudo-randomised such that the first trial of acquisition was always paired and then after all trial types were randomly presented. Conditioning contingencies were counterbalanced, with half of participants receiving the blue square paired with the US and the other half of participants receiving the yellow square paired with the US. The coloured squares were presented for a total of 4000 ms. The aversive sound lasted for 1000 ms, which coterminated with the reinforced CS+’s. Subsequently, a blank screen was presented for 6000 – 8000 ms (see Figure 1).

**Fig 1.**
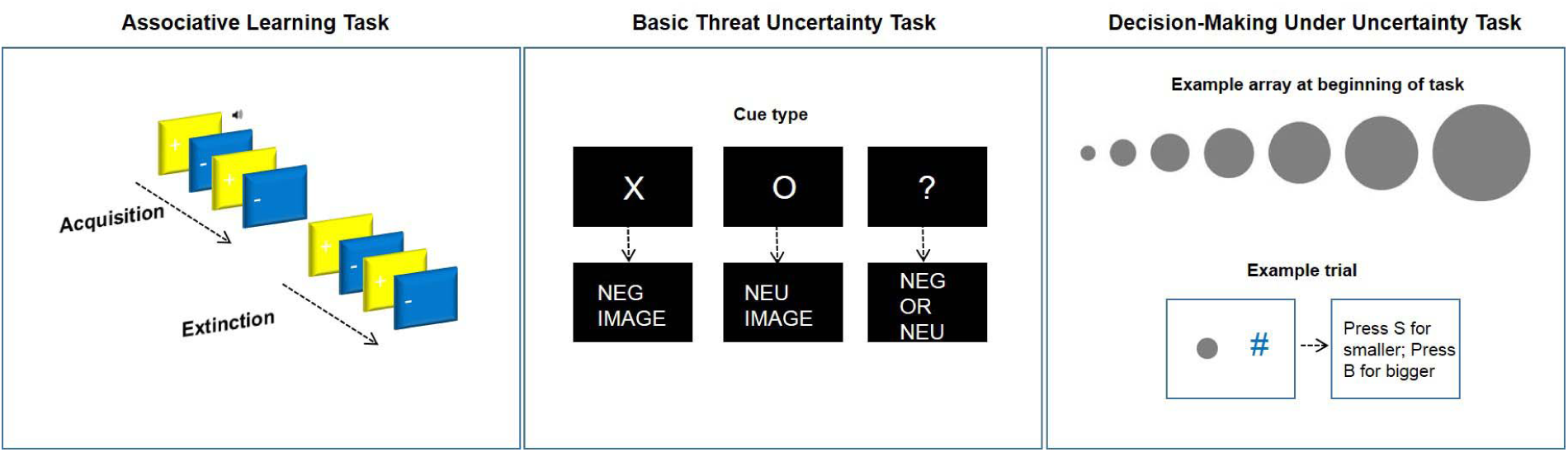
Image displaying the three different uncertainty tasks used in the experiment.

At the end of each block, participants were asked to rate how much they expected the blue square and yellow square to be followed by the sound stimulus, where the scale ranged from 1 (“Don’t Expect”) to 9 (“Do Expect”). Two other 9-point Likert scales were presented at the end of the task. Participants were asked to rate: (1) the valence and (2) arousal of the sound stimulus. The scales ranged from 1 (Valence: very negative; Arousal: calm) to 9 (Valence: very positive; Arousal: excited).

#### 2.3.2 Basic threat uncertainty task

The experimental design for the basic threat uncertainty task was similar or previous work (Gole et al., 2012; Grupe & Nitschke, 2011; Schienle et al., 2010). Participants were required to passively view cues that predicted certain negative, certain neutral or uncertain negative/neutral pictures. Participants were instructed as to which cue predicted a given outcome.

Cues consisted of white courier text with 48 font size (e.g. X, O, ?) presented centrally on a black background. The ‘X’ signified that the participant would receive a certain negative picture. The ‘O’ signified that the participant would receive a certain neutral picture. The ‘?’ signified that the participant could receive a negative or neutral picture.

Thirty-six pictures from the international affective picture system (IAPS) were presented (Lang, Bradley, & Cuthbert, 2005) (for picture list see Table 1). Half of the pictures comprised of negative content (i.e. war, accidents, mutilation, infants in distress) and the other half contained neutral content (i.e. work, children and adults in neutral settings, objects). Based on the original ratings from the IAPS battery, negative pictures were significantly more negative in valence and arousing than neutral pictures, *p*’s < .001. Negative and neutral pictures did not vary in complexity or luminosity, *p*’s > .4. Pictures were presented to fit the 800 × 600 pixel dimensions of the screen.

**Table 1.** Reference numbers to images taken from the International Affective Picture System (IAPS; Lang et al., 2005).

| Negative | Neutral |
| --- | --- |
| 2095 | 2271 |
| 3110 | 2104 |
| 3301 | 9210 |
| 3140 | 2410 |
| 9433 | 7025 |
| 3350 | 2441 |
| 3071 | 2383 |
| 3030 | 7705 |
| 9252 | 2493 |
| 6212 | 7217 |
| 9421 | 2512 |
| 9300 | 2393 |
| 9301 | 2396 |
| 9911 | 7041 |
| 9903 | 9070 |
| 9340 | 7207 |
| 9400 | 2191 |
| 9920 | 5731 |

The task consisted of 36 trials (12 certain negative, 12 certain neutral, 6 uncertain negative, 6 uncertain neutral). Experimental trials were randomised. The pictures were pseudo-randomised such that pictures with similar content would appear in each of the above conditions equally e.g. pictures of war would be shown in both certain negative and uncertain negative conditions. The cue was presented for 4000 ms. The following picture was presented for 4000ms. Lastly, a blank screen was presented for 6000 – 8200 ms (see Figure 1).

Two other 9-point Likert scales were presented at the end of the experiment. Participants were asked to rate: (1) the valence and (2) arousal of the uncertain cue. The scales ranged from 1 (Valence: very negative; Arousal: calm) to 9 (Valence: very positive; Arousal: excited).

#### 2.3.3 Decision-making under uncertainty task

Participants were required to categorise whether a given circle would be larger or smaller than another hypothetical circle. Participants were instructed the following on the computer, ‘In this experiment you will see different size circles. On the left or right you will see a circle. Opposite the circle you will see a #. You will be asked to estimate whether the current circle will likely be bigger or smaller compared to another circle in the array. You will have to wait for 4 seconds before you give your answer using the keyboard.’ Participants then viewed an array of all possible circle sizes for 20 seconds. The array consisted of seven grey circles that ranged from small to large (pixel dimensions: 33 × 33; 63 × 63; 93 × 93; 122 × 123; 152 × 151; 181 × 181; 211 × 211).

The task consisted of 42 trials (6 of each circle size). Experimental trials were randomised. The grey circle and # (pixel dimension: 70 × 81) were counterbalanced to the left and right. The grey circle and # were presented for 4000ms. The response slide was presented for 2000ms and asked participants to ‘Press S for smaller; Press B for bigger’. Lastly, a fixation screen was presented for 6000 – 7000 ms (see Figure 1).

### 2.4 Questionnaires

STAI (Spielberger, Gorsuch, Lushene, Vagg, & Jacobs, 1983) and IU questionnaires (Freeston, Rhéaume, Letarte, Dugas, & Ladouceur, 1994) were measured.

### 2.5 Behavioural data scoring

Rating data were reduced for each participant by calculating their average responses for each experimental condition using the E-Data Aid tool in E-Prime (Psychology Software Tools Ltd, Pittsburgh, PA).

#### 2.5.1 Associative threat learning task

Average expectancy ratings were calculated for the following conditions (Acquisition CS+, Acquisition CS-, Extinction Early CS+, Extinction Early CS-, Extinction Late CS+, Extinction Late CS-).

#### 2.5.2 Decision-making under uncertainty task

Reaction times under 200ms were discarded. Remaining reaction times greater or equal to 200ms were z--scored to control for interindividual differences in reaction time. The reaction times were averaged for each trial type (Circles 1-7, from small to large) and then collapsed over the following conditions: Certain (Circles 1 and 7), Mildly Uncertain (Circles 2 and 6), Quite Uncertain (Circles 3 and 5) and Very Uncertain (Circle 4).

Choices were coded into 1 and 0, stimuli that were predicted as bigger were assigned a value of 1 and stimuli that were predicted as smaller were assigned a value of 0. Average values for each condition (Circles 1-7, from small to large) essentially acted as subjects’ probability of stimulus size. For example, a value of 1, meant that the subject always picked bigger, whilst a value of .5 meant that the subject picked between bigger and smaller equally.

### 2.6 Physiological acquisition

Physiological recordings were obtained using AD Instruments (AD Instruments Ltd, Chalgrove, Oxfordshire) hardware and software.

Electrodermal activity was measured with dry MLT116F silver/silver chloride bipolar finger electrodes that were attached to the distal phalanges of the index and middle fingers of the non-dominant hand. A low constant-voltage AC excitation of 22 mV_rms_ at 75 Hz was passed through the electrodes, which were connected to a ML116 GSR Amp, and converted to DC before being digitized and stored. An ML138 Bio Amp connected to an ML870 PowerLab Unit Model 8/30 amplified the skin conductance signal, which was digitized through a 16-bit A/D converter at 1000 Hz.

Facial EMG measurements of the corrugator supercilii muscles were obtained by using two pairs of 4 mm Ag/AgCl bipolar surface electrodes connected to the ML138 Bio Amp. The centres of each pair of bipolar surface electrodes were approximately 15 mm apart. The reference electrode was a singular 8mm Ag/AgCl electrode, placed upon the middle of the forehead, and connected to the ML138 Bio Amp. Before placing the EMG sensors the skin site was cleaned with distilled water and slightly abraded with isopropyl alcohol skin prep pads, to reduce skin impedance to an acceptable level (below 20kΩ).

### 2.7 Physiological scoring

The physiological parameters were extracted using AD Instruments software.

#### 2.7.1 Associative threat learning task

CS+ unpaired and CS-trials were included in the analysis, but CS+ paired trials were discarded to avoid sound confounds. Skin conductance responses (SCR) were scored when there was an increase of skin conductance level exceeding 0.03 microSiemens (Dawson et al., 2000). The amplitude of each response was scored as the difference between the onset and the offset (maximum deflection prior to the signal flattening out or decreasing). SCR onsets and offsets were counted if the SCR onset was within 0.5-3.5 seconds following CS onset (Morriss, Chapman, Tomlinson, & Van Reekum, 2018). Trials with no discernible SCRs were scored as zero. SCR magnitudes were square root transformed to reduce skew and were z-scored to control for interindividual differences in skin conductance responsiveness (Ben Shakhar, 1985). SCR magnitudes were calculated from remaining trials by averaging SCR transformed values and zeros for each condition (Acquisition CS+, Acquisition CS-, Extinction Early CS+, Extinction Early CS-, Extinction Late CS+, Extinction Late CS-).

A high-pass filter at 20hz was applied to the raw corrugator online (Solnik, DeVita, Rider, Long, & Hortobágyi, 2008). The corrugator signal was root mean squared offline (Fridlund & Cacioppo, 1986). Baseline corrected second by second means over the course of the CS (4 seconds) were extracted for corrugator supercilii. Baseline mean values were taken from the 2 second period before each trial began. The corrugator supercilii was extracted for each condition (Acquisition CS+, Acquisition CS-, Extinction Early CS+, Extinction Early CS-, Extinction Late CS+, Extinction Late CS-).

#### 2.7.2 Basic threat uncertainty task

As above, the same criteria were applied to SCR and EMG during the basic threat uncertainty task. SCR onsets and offsets were counted if the SCR onset was within 0.5-3.5 seconds following cue and picture onset. SCR magnitudes were calculated from remaining trials by averaging SCR transformed values and zeros for each condition (Cue Certain Negative, Cue Certain Neutral, Cue Uncertain, Certain Negative Picture, Certain Neutral Picture Uncertain Negative Picture, Uncertain Neutral Picture).

Baseline corrected second by second means over the course of the cue and picture (4 seconds) were extracted for corrugator supercilii. Baseline mean values were taken from the 2 second period before each cue or picture began. The corrugator supercilii was extracted for each condition (Cue Certain Negative, Cue Certain Neutral, Cue Uncertain, Certain Negative Picture, Certain Neutral Picture Uncertain Negative Picture, Uncertain Neutral Picture).

#### 2.7.3 Decision-making under uncertainty task

The same criteria were applied to SCR and EMG during the decision-making under uncertainty task. SCR onsets and offsets were counted if the SCR onset was within 0.5-3.5 seconds following the decision array. SCR magnitudes were calculated from remaining trials by averaging SCR transformed values and zeros for each condition (Circles 1-7, from small to large).

Baseline corrected second by second means over the course of the decision array (4 seconds) were extracted for corrugator supercilii. Baseline mean values were taken from the 2 second period before each decision array began. The corrugator supercilii was extracted for each condition (Circles 1-7. from small to large).

For SCR and corrugator supercilii, the trial types were collapsed into: Certain (Circles 1 and 7), Mildly Uncertain (Circles 2 and 6), Quite Uncertain (Circles 3 and 5) and Very Uncertain (Circle 4).

### 2.8 Behaviour and physiology analysis

The analyses were conducted using the mixed procedure in SPSS 21.0 (SPSS, Inc; Chicago, Illinois). For all models a diagonal covariance matrix was used for level 1. Random effects included a random intercept for each individual subject, where a variance components covariance structure was used. A maximum likelihood estimator and the least significance difference procedure for pairwise comparisons was used for the multilevel models. The following individual difference predictor variables were entered into all of the multilevel models: IU and STAI.

The specificity of IU with respect to STAI is reported when a significant interaction was observed i.e. IU with Stimulus, Time or Second. Then, follow-up pairwise comparisons are performed on the estimated marginal means, adjusted for the predictor variables (IU, STAI). Any interaction with IU was followed up with pairwise comparisons of the means between the conditions for IU estimated at the specific values of + or - 1 SD of mean IU (Morriss, Macdonald & van Reekum, 2016; Morriss, McSorely & van Reekum, 2017). These data are estimated from the multilevel model of the entire sample, not unlike performing a simple slopes analysis in a multiple regression analysis.

#### 2.8.1 Associative threat learning task

Separate multilevel models were conducted on expectancy ratings and SCR magnitude for each phase (Acquisition, Extinction). For expectancy ratings and SCR magnitude during the acquisition phase, Stimulus (CS+, CS-) was entered at level 1 and individual subjects at level 2. For expectancy ratings and SCR magnitude during the extinction phase, Stimulus (CS+, CS-) and Time (Early, Late) were entered at level 1 and individual subjects at level 2. The models for the corrugator supercilii in acquisition included Stimulus (CS+, CS-) and Second (1,2,3,4) at level 1 and individual subjects at level 2. In addition, the model for corrugator supercilii in extinction included Stimulus (CS+, CS-), Time (Early, Late) and Second (1,2,3,4) at level 1 and individual subjects at level 2.

#### 2.8.2 Basic threat uncertainty task

SCR magnitude to the cue during the basic threat uncertainty task was assessed by including Cue (Certain Negative, Certain Neutral and Uncertain) at level 1 and individual subjects at level 2. To examine the impact of cue on SCR magnitude to the picture, an additional model was conducted, where Cue (Certain, Uncertain) and Picture (Negative, Neutral) was entered at level 1 and individual subjects at level 2. Furthermore, corrugator supercilii activity to the cue was assessed by including Cue (Certain Negative, Certain Neutral and Uncertain) and Second (1,2,3,4) at level 1 and individual subjects at level 2. Lastly, to assess the impact of cue on corrugator supercilii activity during the picture, another model was conducted where Cue (Certain, Uncertain), Picture (Negative, Neutral) and Second (1,2,3,4) were entered at level 1 and individual subjects at level 2.

#### 2.8.3 Decision-making under uncertainty task

Choices to the decision array were assessed by including Levels of Uncertainty (Circles 1-7, from small to large) at level 1 and individual subjects at level 2.

Reaction time and SCR magnitude to the decision array was assessed by including Levels of Uncertainty (Certain, Mildly Uncertain, Quite Uncertain and Very Uncertain) at level 1 and individual subjects at level 2. Furthermore, corrugator supercilii activity to the decision array was assessed by including Levels of Uncertainty (Certain, Mildly Uncertain, Quite Uncertain and Very Uncertain) and Second (1,2,3,4) at level 1 and individual subjects at level 2.

## 3. Results

### 3.1 Questionnaires

Similar distributions and internal reliability of scores were found for the self-report anxiety measures, STAI (*M* = 45.18; *SD* = 9.09; range = 26-66; *α* = .88), IU (*M* = 61.82; *SD* = 18.02; range = 33-103; *α* = .93). There was a significant positive correlation between IU and STAI, *r*(43) = .475, *p* = .001.

### 3.2 Associative threat learning task

#### 3.2.1 Ratings

Participants rated the sound stimulus as aversive (*M* = 2.14, *SD* = 1.3, range 1-6, where 1 = very negative and 9 = very positive) and arousing (*M* = 7.19, *SD* = 1.4, range 3-9 where 1 = calm and 9 = excited).

For the expectancy ratings, during acquisition participants reported greater expectancy of the sound with the CS+, compared to CS-[Stimulus: *F*(1, 79.747) = 119.416, *p* < .001] (for descriptive statistics see Table 2). No other significant interactions with IU group or STAI were found for the ratings during acquisition, max *F* =.730.

**Table 2.** Associative threat learning task summary of means (SD) for each dependent measure as a function of stimulus (CS+ and CS-), separately for acquisition, early extinction and late extinction.

| Measure | Acquisition |  | Early Extinction |  | Late Extinction |  |
| --- | --- | --- | --- | --- | --- | --- |
|  | CS+ | CS- | CS+ | CS- | CS+ | CS- |
| Square root transformed and z-scored SCR magnitude ($\sqrt{\mu\text{S}}$) | .38<br>(.42) | .05<br>(.34) | .07<br>(.37) | -.23<br>(.28) | -.05<br>(.39) | -.15<br>(.37) |
| Corrugator supercilii activity ($\mu\text{V}$) | .76<br>(1.63) | .29<br>(.70) | .49<br>(1.42) | .25<br>(.87) | .17<br>(.75) | .14<br>(.56) |
| Expectancy rating (1-9) | 6.96<br>(1.68) | 2.50<br>(2.12) | 4.64<br>(2.11) | 2.23<br>(2.26) | 3.45<br>(2.06) | 2.42<br>(2.46) |

Note: SCR magnitude ( $\sqrt{\mu\text{S}}$ ), square root transformed and z-scored skin conductance magnitude measured in microSiemens. Baseline corrected corrugator supercilii activity ( $\mu\text{V}$ ), measured in microVolts. Expectancy ratings from 1-9, where 1 is don't expect and 9 is very much expect.

During extinction, participants reported greater expectancy of the sound with the CS+, compared to CS-[Stimulus: *F*(1, 121.742) = 61.407, *p* < .001]. The expectancy ratings dropped over time [Time: *F*(1, 121.742) = 5.274, *p* = .023; Stimulus x Time: *F*(1, 121.742) = 9.878, *p* = .002]. Follow-up pairwise comparisons revealed that the expectancy rating of the sound with the CS+ dropped significantly from early to late extinction, *p* < .001. However, the expectancy rating of the CS-with the sound remained low and did not change with time, *p* = .546. No other significant interactions with IU group or STAI were found for the ratings during extinction, max F = .810.

#### 3.2.2 SCR

As expected, CS+ stimuli elicited larger SCR magnitudes than CS-during acquisition [Stimulus: *F*(1,79.516) = 16.134, *p* < .001] (see, Table 2). There were no significant interactions between Stimulus x IU or STAI for SCR magnitude during acquisition, max *F* = 1.092.

During extinction, SCR magnitude was greater for the CS+ vs. CS-[Stimulus: *F*(1,160.257) = 14.186, *p* <.001] (see Table 2). Additionally, SCR magnitude was greater for the CS+ vs. CS-during early extinction, p < .001, but not for late extinction, p =.224 [Stimulus x Time: *F*(1, 160.257) = 4.253, *p* =. 041].

As predicted IU was related to SCR magnitude during extinction [Stimulus x Time x IU interaction: *F*(1,160.257) = 6.760, *p* =.010]. Further inspection of follow-up pairwise comparisons for early vs. late extinction at IU ±1 SD from the mean on the regression line showed lower IU (1 SD below the mean) to be associated with significantly greater SCR magnitude in early extinction to the CS+, relative to the CS-, *p* < .001, but not in late extinction, *p* = .313 (see, Figure. 2). In contrast, higher IU (1 SD above the mean) was associated with greater SCR magnitude to CS+ vs. CS-in both early extinction, *p* = .05, and late extinction, *p* = .011. Furthermore, low IU was associated with reduced SCR magnitude to the CS+ from early to late extinction, *p* = .012, and an increase in SCR magnitude to the CS-from early to late extinction, *p* = .047.^1^ No other significant main effects or interactions were found with Stimulus, Time, IU or STAI for SCR magnitude during extinction, max F = 1.919.

**Fig 2.**
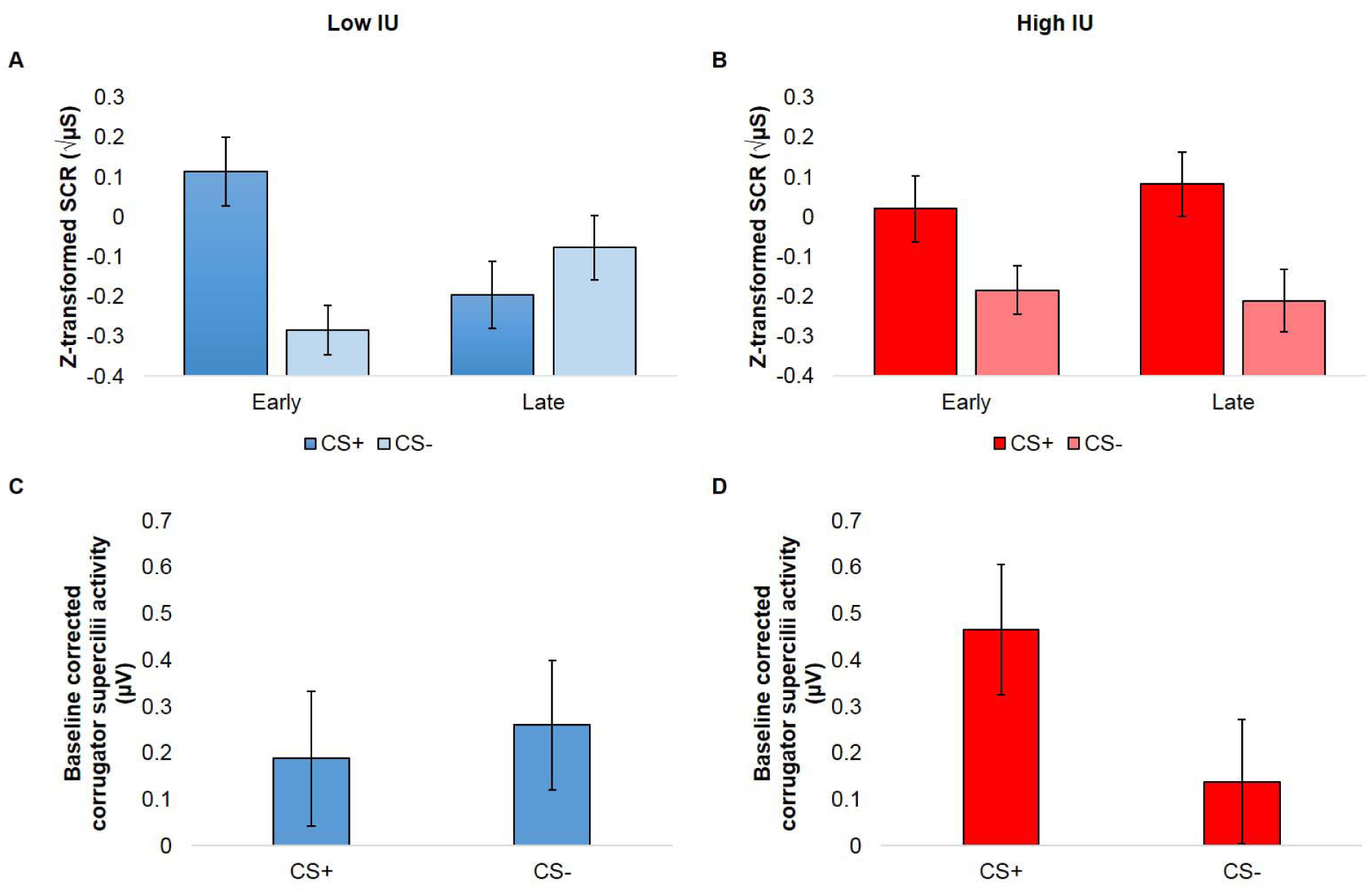
Bar graphs depicting IU estimated at + or - 1 SD of mean IU (controlling for STAI) from the multilevel model analysis for SCR magnitude and corrugator supercilii activity during extinction phase of the associative threat learning task. High IU, relative to low IU individuals were found to show heightened SCR magnitude and corrugator supercilii responding to the CS+ versus CS-cue during extinction. Bars represent standard error at + or – 1 SD of mean IU. Square root transformed and z-scored SCR magnitude (μS), skin conductance magnitude measured in microSiemens. Baseline corrected corrugator supercilii activity (μV), measured in microVolts.

#### 3.2.3 Corrugator supercilii activity

During acquisition larger corrugator supercilii activity was observed to the CS+ vs. CS-[Stimulus: *F*(1,167.048) = 19.865, *p* < .001] (see, Table 2). There were no significant interactions between Stimulus or Second x IU or STAI for corrugator supercilii activity during acquisition, max *F* = 2.202.

During extinction larger corrugator supercilii activity was observed to the CS+ vs. CS-at trend [Stimulus: *F*(1,398.194) = 3.356, *p* = .068] (see, Table 2). Corrugator supercilii activity reduced across early to late extinction [Time: *F*(1,398.194) = 9.256, *p* = .003]. As expected, individuals scoring higher in IU were shown to exhibit larger larger corrugator supercilii activity to the CS+ vs. CS-during extinction, *p* = .02, compared to individuals scoring lower in IU, *p* = .499 [Stimulus x IU: *F*(1,398.194) = 6.438, *p* = .012] (see Figure 2). No other significant interactions between Stimulus or Second x IU or STAI were observed for corrugator supercilii activity during extinction, max *F* = 2.911.

### 3.3 Basic threat uncertainty task

#### 3.3.2 SCR

Certain negative cues elicited larger SCR magnitudes, compared to the certain neutral cue and uncertain cue, p’s < .001 [Cue: F(1,79.516) = 16.134, p < .001] (see, Table 3). However, there was no significant difference between SCR magnitudes for certain neutral cues and uncertain cues, p = .334. There were no significant interactions between Cue x IU or STAI for SCR magnitude, max *F* = .685.

As expected negative pictures elicited larger SCR magnitudes, compared to neutral pictures [Picture: F(1,151.603) = 13.369, p < .001].^2^ No other significant interactions between Cue x Picture x IU or STAI for SCR magnitude were observed, max *F* = 3.543.

**Table 3.** Basic threat uncertainty task summary of means (SD) for each dependent measure as a function of the cue and picture period separately.

| Measure | Certain Negative Cue |  | Certain Neutral Cue |  | Uncertain Cue |
| --- | --- | --- | --- | --- | --- |
| Square root transformed and z-scored SCR magnitude ($\sqrt{\mu\text{S}}$) | .14 (.26) | | -.10 (.22) | | -.15 (.28) |
| Corrugator supercilii activity ($\mu\text{V}$) | -.06 (.81) | | -.03 (.45) | | .01 (.46) |
|  | Certain Cue |  | Uncertain Cue |  |  |
|  | Negative | Neutral | Negative | Neutral |  |
| Square root transformed and z-scored SCR magnitude ($\sqrt{\mu\text{S}}$) | .06 (.35) | -.14 (.23) | .14 (.45) | -.05 (.40) | |
| Corrugator supercilii activity ($\mu\text{V}$) | .89 (1.22) | -.04 (.63) | .74 (1.20) | .05 (.74) | |
| Note: SCR magnitude ($\sqrt{\mu\text{S}}$), square root transformed and z-scored skin conductance magnitude measured in microSiemens. Baseline corrected corrugator supercilii activity ($\mu\text{V}$), measured in microVolts. | | | | | |

#### 3.3.3 Corrugator supercilii activity

No significant main effects or interactions between Cue x IU or STAI for corrugator supercilii activity were observed, max *F* = 1.242. Larger corrugator supercilii activity was found to negative, compared to neutral pictures [Picture: *F*(1,397.444) = 119.813, *p* < .001; Picture x Second: *F*(1,397.444) = 11.742, *p* < .001] (see Table 3).

An interaction between Cue, Picture and STAI emerged for corrugator supercilii activity [Cue x Picture x STAI interaction: *F*(1, 397.444) = 14.956, *p* <.001] (see Figure 3). Follow-up pairwise comparisons revealed that higher STAI was associated with larger corrugator supercilii activity to both certain negative pictures, compared to certain neutral pictures, and uncertain negative pictures, compared to uncertain neutral pictures, *p*’s < .001. Moreover, higher STAI was associated with greater corrugator supercilii activity to certain neutral pictures, compared to uncertain neutral pictures, *p* < .001, but there was no significant difference between certain negative pictures, compared to uncertain negative pictures, *p* = .383. Lower STAI was associated with larger corrugator supercilii activity to certain negative pictures, compared to certain neutral pictures, *p* < .001, but not to uncertain negative, compared to uncertain neutral pictures, *p* =.252. In addition, lower STAI was associated with greater corrugator supercilii activity to certain pictures versus uncertain pictures, regardless of valence, *p*’s < .015.

**Fig 3.**
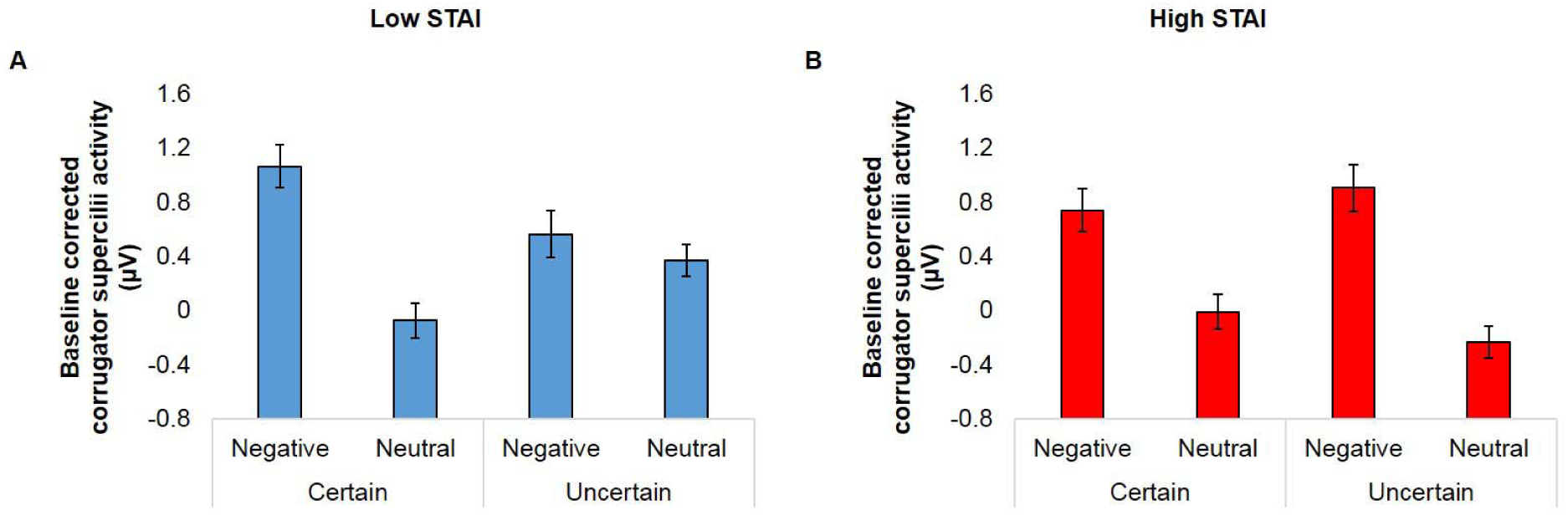
Bar graphs depicting STAI estimated at + or - 1 SD of mean STAI (controlling for IU) from the multilevel model analysis for corrugator supercilii activity during the basic uncertainty threat task. High STAI, relative to low STAI individuals were found to show larger corrugator supercilii activity to both certain and uncertain negative pictures, relative to certain and uncertain neutral pictures. Low STAI, relative to high STAI individuals were only found to show larger supercilii activity to certain negative versus certain neutral pictures, and not uncertain negative versus uncertain neutral pictures. Bars represent standard error at + or – 1 SD of mean STAI. Baseline corrected corrugator supercilii activity (μV), measured in microVolts.

No other significant interactions between Cue x Picture x IU or STAI for corrugator supercilii activity were observed, max *F* = 3.514.

### 3.4 Decision-making under uncertainty task

Three subjects missed responses to over half of the conditions and therefore were excluded from the analysis of the decision-making task.

#### 3.4.1 Behavioural responses

Subjects predicted circle size as expected, such that there were significant differences in choices based on probability, particularly for circles in the middle (i.e. 2, 3,4,5), p’s <.05. Choices based on probability did not differ between circles 1 and 2, and circles 5, 6 and 7, p’s > .05 [Levels of Uncertainty: *F*(1, 61.359) = 33.646, *p* <.001] (see table 4). For choices during the decision-making task there was a trend between Levels of Uncertainty and IU [Levels of Uncertainty x IU interaction: *F*(1, 53.654) = 2.147, *p* =.054]. Follow-up pairwise comparisons revealed that higher IU, relative to lower IU was associated with more accurate choice predictions of whether a circle would be likely bigger or smaller than the another circle shown in the array. For example, high IU was associated with better probabilistic choices for circles in the middle (i.e. 2,3,4) p’s < .05, whilst low IU was not, p’s > .2 (see Figure 4).

**Table 4.** Decision-making under uncertainty task summary of means (SD) for each dependent measure as a function of stimulus (Circles 1-7, from small to large) and levels of uncertainty (Certain, Mildy Uncertain, Quite Uncertain, Very Uncertain) separately.

| Measure | 1 | 2 | 3 | 4 | 5 | 6 | 7 |
| --- | --- | --- | --- | --- | --- | --- | --- |
| Choice | .83<br>(.31) | .68<br>(.32) | .48<br>(.33) | .30<br>(.31) | .18<br>(.25) | .14<br>(.26) | .12<br>(.27) |

| Measure | Certain | Mildy<br>Uncertain | Quite<br>Uncertain | Very Uncertain |
| --- | --- | --- | --- | --- |
| Square root transformed and z-scored SCR magnitude ($\sqrt{\mu S}$) | -.007 (.22) | -.009 (.13) | -.03 (.23) | .03 (.37) |
| Corrugator supercilii activity ($\mu V$) | -.16 (.64) | -.18 (.74) | -.001 (.72) | -.01 (.77) |
| Z-scored reaction time (ms) | .009 (.27) | -.01 (.16) | -.02 (.25) | .06 (.42) |

Note: Choices were coded into 1 and 0, stimuli that were predicted as bigger were assigned a value of 1 and stimuli that were predicted as smaller were assigned a value of 0. Average values for each condition (Circles 1-7, from small to large) acted as subjects' probability of stimulus size. For example, a value of 1, meant that the subject always picked bigger. SCR magnitude ( $\sqrt{\mu S}$ ), square root transformed and z-scored skin conductance magnitude measured in microSiemens. Baseline corrected corrugator supercilii activity ( $\mu V$ ), measured in microVolts. Reaction times (ms), z-scored reaction times measured in milliseconds.

**Fig. 4.**
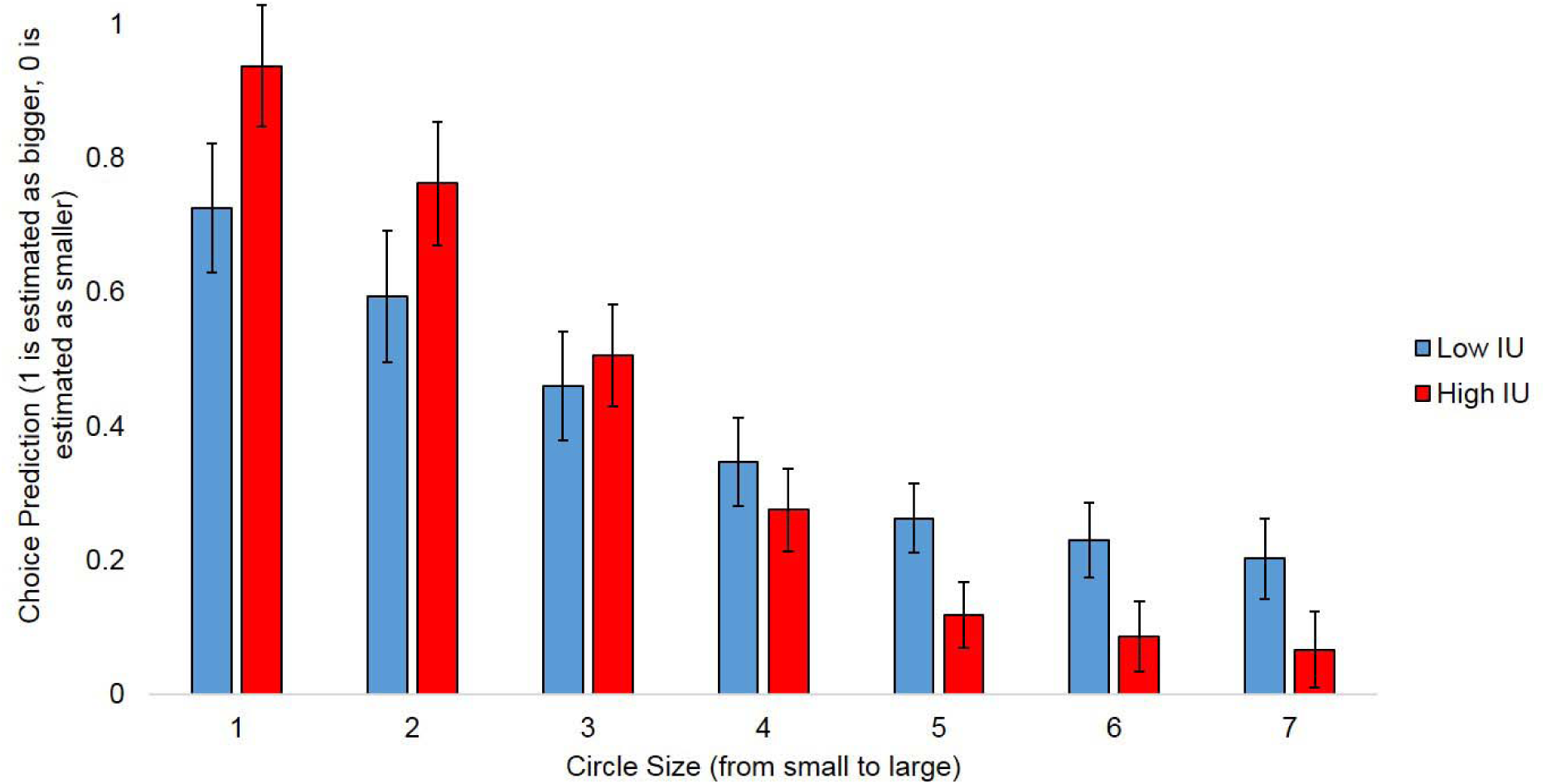
Graph illustrating IU estimated at + or - 1 SD of mean IU (controlling for STAI) from the multilevel model analysis for choices made during the decision-making under uncertainty task. Higher IU relative to lower IU was associated with more accurate choice predictions of whether a circle would be likely bigger or smaller than another circle shown in the array. This effect was particularly evident at the most certain circles (1 and 7). Choices were coded into 1 and 0, stimuli that were predicted as bigger were assigned a value of 1 and stimuli that were predicted as smaller were assigned a value of 0. Average values for each condition (Circles 1-7, from small to large) acted as subjects’ probability of stimulus size.

No other significant main effects or interactions between Levels of Uncertainty x IU or STAI for choices or reaction times were observed, max *F* = .314.

#### 3.4.2 SCR

For SCR magnitude an interaction at trend between Levels of Uncertainty and IU emerged [Levels of Uncertainty x IU interaction: *F*(1, 42) = 2.369, *p* =.068] (see Figure 5). Follow-up pairwise comparisons revealed that lower IU was associated with greater SCR magnitude to very uncertain versus certain stimuli, *p* = .020. No other significant main effects or interactions between Levels of Uncertainty x STAI for SCR magnitudes were observed, max *F* = .798.

**Fig 5.**
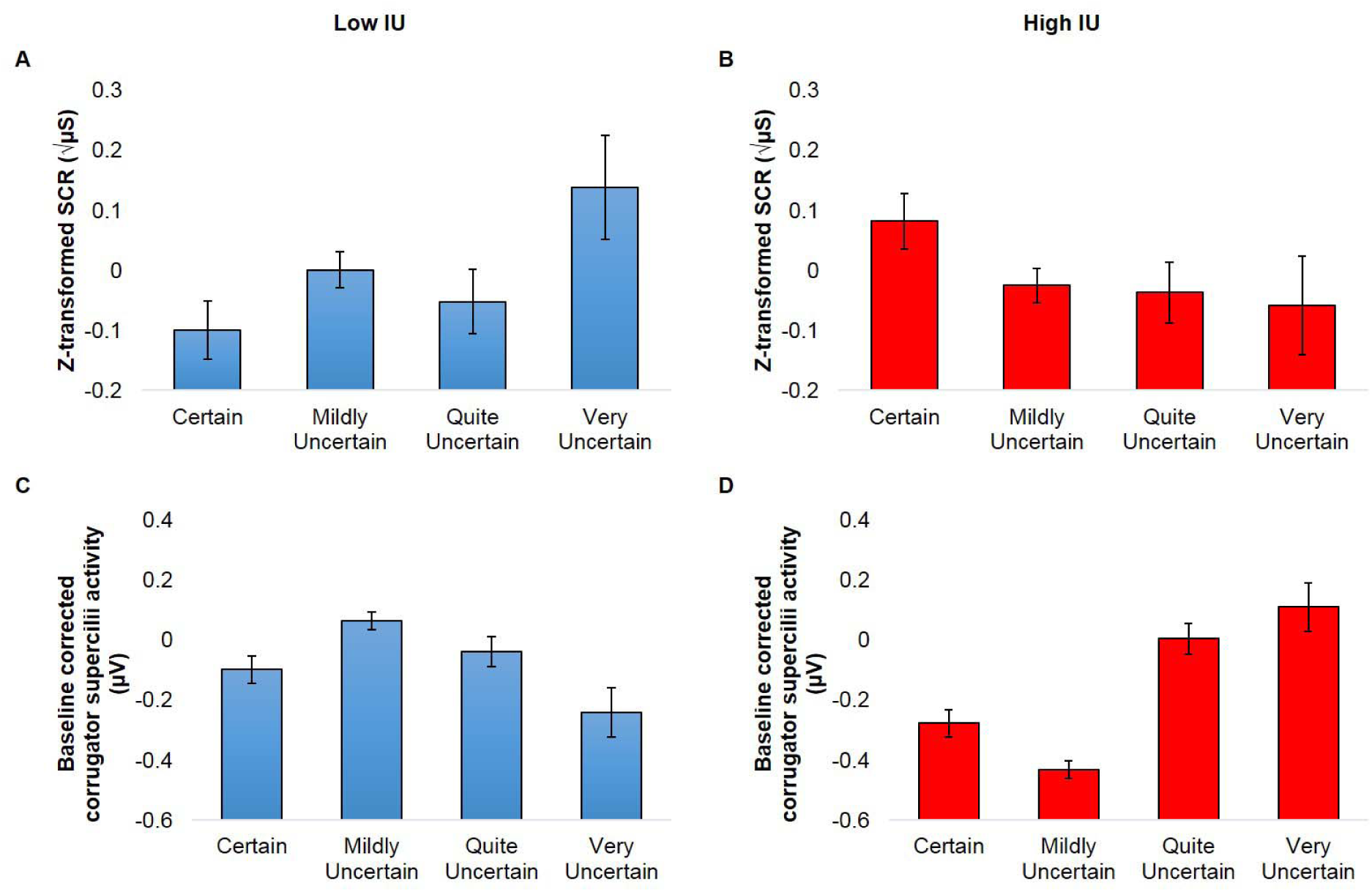
Bar graphs showing IU estimated at + or - 1 SD of mean IU (controlling for STAI) from the multilevel model analysis for SCR magnitude and corrugator supercilii activity during the decision-making under uncertainty task. High IU, relative to low IU individuals were found to show reduced corrugator supercilii activity to the certain and mildy uncertain conditions, compared to the quite and very uncertain conditions. Low IU individuals displayed larger SCR magnitude to the very uncertain condition, compared to the certain condition, whilst high IU individuals did not show any significant differences in SCR magnitude across conditions. Bars represent standard error at + or – 1 SD of mean IU. Square root transformed and z-scored SCR magnitude (μS), skin conductance magnitude measured in microSiemens. Baseline corrected corrugator supercilii activity (μV), measured in microVolts.

#### 3.4.3 Corrugator supercilii activity

IU was related to corrugator supercilii activity during the decision-making task [Levels of Uncertainty x IU interaction: *F*(1,193.140) = 6.830, *p* <.001] (see Figure 4). Further inspection of follow-up pairwise comparisons revealed that higher IU was associated with significantly reduced corrugator supercilii activity to certain and mildy uncertain, relative to quite and very uncertain stimuli, *p*’s < .032. Lower IU was associated only with larger corrugator activity to the mildy uncertain relative to the very uncertain, *p* = .024.^3^ No other significant interactions between Levels of Uncertainty x IU or STAI for the corrugator supercilii were observed, max *F* = 1.950.

## 4. Discussion

Here anticipatory physiological responses to uncertainty in three different contexts (associative threat learning, basic threat uncertainty, decision-making) within the same sample were examined. IU made different contributions to anticipatory physiological responses during each task. IU was found to modulate both SCR magnitude and corrugator supercilii activity for the associative threat learning and decision-making contexts. However, STAI was found to modulate corrugator supercilii activity during the basic threat uncertainty context. Taken together this research helps us further dissect the role of IU during different uncertain contexts.

For the associative threat learning task, typical patterns of threat acquisition were observed, such that larger SCR magnitude, corrugator supercilii activity and expectancy ratings were found for the learned threat vs. safety cues (for review see, Lonsdorf et al., 2017). In addition, effects of threat extinction were observed as larger SCR magnitude and expectancy ratings were found for the early part of the extinction phase, relative to the late part of extinction phase. As expected, higher IU was associated with reduced threat extinction, as shown by larger SCR magnitude and corrugator supercilii activity to learned threat vs. safety cues across the extinction phase. The observed IU-related effects on SCR magnitude and corrugator supercilii activity during extinction were specific to IU, over STAI. These results replicate prior work which has shown that high IU is associated with poorer extinction outcomes (Dunsmoor et al., 2015; Lucas et al., 2018; Morriss et al., 2015; Morriss, Christakou, et al., 2016; Morriss, Macdonald, et al., 2016). Overall, the findings from the current study provide further evidence that threat extinction may induce uncertainty-related anxiety in individuals who score higher in IU because threat extinction is inherently uncertain, as the contingencies are unknown.

In the basic threat uncertainty task, similar patterns of physiological responses to previous research were observed (Grupe et al., 2011). For example, larger SCR magnitude was found to the certain negative cue, relative to the certain neutral and uncertain cues. Additionally, larger SCR magnitude and corrugator supercilii activity was observed for negative, compared to neutral pictures (Lang & Bradley, 2010; Lang, Greenwald, Bradley, & Hamm, 1993). An interaction between the cue, picture and STAI revealed that higher STAI was associated with larger corrugator supercilii activity to negative images that followed an uncertain and certain cue, whilst lower STAI was associated with only larger corrugator supercilii activity to certain versus uncertain negative images. Importantly, the observed STAI-related effects on corrugator supercilii activity during the basic threat uncertainty task were specific to STAI, over IU. Although these results were not expected, the results aren’t surprising given the lack of reported IU effects on physiological measures during these type of tasks (Bennett et al., 2018; Grupe & Nitschke, 2011). There may be multiple reasons for this result: (1) the task is not that uncertain given the known contingencies and therefore is not as motivationally relevant to individuals who score high in IU, and (2) the task presents many different types of negative images, tapping into broader negative affective states i.e. disgust, sadness, and fear, which may be more motivationally relevant for STAI, given that the items are relevant for both anxiety and depression (Grös, Antony, Simms, & McCabe, 2007).

For the decision-making under uncertainty task, all participants gave the typical choices when predicting whether a hypothetical circle would likely be bigger or smaller than the one that was presented. For example, if the circle presented was small than participants were more likely to pick that the circle under the hash sign would likely be bigger. Notably, high IU, relative to low IU was associated with better choice accuracy (e.g. more accurate probabilities). This result may be interpreted to reflect differences in motivational relevance for high and low IU individuals. High IU individuals may have been more engaged in the task in order to reduce uncertainty. In addition, high IU individuals relative to low IU individuals exhibited reduced corrugator supercilii activity during the anticipation of making a decision to certain stimuli, compared to uncertain stimuli. This can be interpreted as High IU individuals expressing relief during certain stimuli and expressing distress during uncertain stimuli. Lastly, lower IU was associated with larger SCR magnitudes to the uncertain stimuli, relative to certain stimuli. Whilst this effect for SCR magnitude was in the opposite direction for higher IU, albeit not significant. These results are difficult to interpret, as heightened arousal to uncertainty would be expected for high IU. However, given the corrugator supercilii results with IU during the decision-making task, these changes in SCR magnitude may reflect arousal from liking uncertain stimuli in low IU individuals. Additional research is needed to clarify the extent to which arousal in this decision-making task is related to liking/distress.

Taken together the results from the decision-making task are in line with previous work related to IU, showing that high IU individuals may seek to reduce uncertainty (Jacoby et al., 2014; Jacoby et al., 2016; Ladouceur et al., 1997), and may feel relief from certainty. Previous work has shown individual differences in IU to modulate decision-making during tasks with valenced outcomes (Carleton et al., 2016; Luhmann et al., 2011; Tanovic, Hajcak, et al., 2018; Tanovic, Pruessner, & Joormann, 2018). Here it was shown that individual differences in IU modulate decision-making in the absence of valenced outcomes or consequences, thus suggesting that anticipating making a decision under uncertainty is enough to induce heightened physiological responses. Notably, during this task no feedback was given, which may have ramped up the uncertainty for high IU individuals. It will be important in future research to examine decision-making under uncertainty with feedback, as well as different valenced outcomes, in order to examine whether these parameters change IU-related physiological profiles.

The preliminary results from the current study provide us with a more nuanced understanding of IU’s role on anticipatory physiological responses during different uncertain contexts. IU may be more relevant for modulating anticipatory physiological responses in situations: (1) where the contingencies are unknown such as threat extinction, and (2) where decisions under uncertainty are made in the absence of immediate outcomes/feedback. IU may not be as relevant for modulating anticipatory physiological responses in situations where the rules of uncertainty are known, as in basic threat uncertainty tasks. In general, from these results it can be speculated that situations with unknown uncertainties may be more relevant for IU-based anticipatory physiological responses. This notion of the unknown as the ultimate stressor for IU is in line with modern definitions, where IU has been described as a ‘dispositional incapacity to endure the aversive response triggered by the perceived absence of salient, key, or sufficient information, and sustained by the associated perception of uncertainty’ (Carleton, 2016b, p 31). Perhaps, another potential avenue of research in IU is to examine not only the parameters of certainty-uncertainty but also known-unknown (see Fig 6).

**Fig 6.**
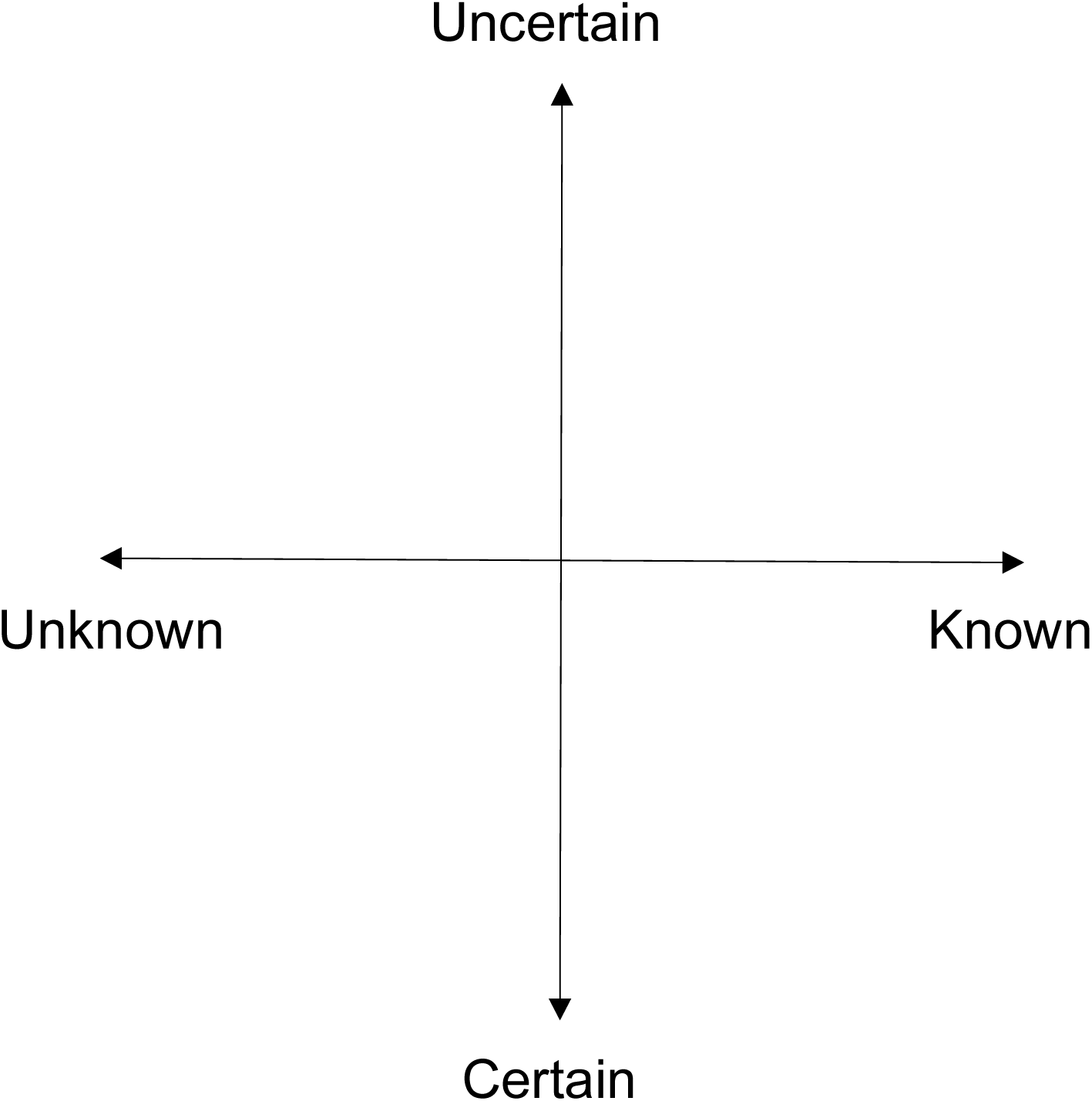
Image displaying uncertainty-certainty and unknown-known parameters together. Future research is needed on how these different parameters impact IU-related phenomena.

In order to assess the generalisability of the physiological profiles observed in this study, it will be important to examine the role of IU on anticipatory responses using other measures (i.e. behavioural, physiological and neural) and samples. In addition, further work is needed to examine how different levels of uncertainty/unknowns in the absence and presence of valenced outcomes across contexts modulate these IU-related physiological profiles (Shihata et al., 2016). Such research will be important because it may reveal underlying mechanisms that serve as common denominators across IU-related psychopathology. This is in line with the Research Domain Criteria (Insel et al., 2010), which aims to identify dimensions of observable behaviours that underpin mental health problems across diagnostic boundaries. Ultimately, this body of research will feed into developing future transdiagnostic models of IU.

The current study had a number of shortcomings. The uncertain contexts that were examined were based on the most popular areas of research on uncertainty (Morriss, Gell & van Reekum, 2019). There may have been other uncertain contexts that would be relevant to address and which have been missed here (i.e. attentional inhibition, perception). Furthermore, there may have been more elegant ways to subdivide the different uncertain contexts (e.g. uncertainty/unknowns and aversion/reward outcomes). There are other psychophysiological measures that should be examined in relation to IU and different uncertain contexts i.e. startle, heart rate variability. The experiment was conducted on a relatively small sample from the UK as a proof of concept. To assess the robustness of these effects future research should aim to use larger and more diverse samples from around the world.

In conclusion, these initial results provide insight into how IU is related to anticipatory physiological responses during different types of uncertainty, which will be relevant for understanding uncertainty-induced anxiety and potential treatment targets (Carleton, 2016a, 2016b; Grupe & Nitschke, 2013). Further research is needed to explore how individual differences in IU modulate anticipatory physiological responses during contexts with varying levels of uncertainty/unknowns and valences outcomes.

## Author Note

This research was supported by a British Academy Small Research Grant (SG163121) awarded to Jayne Morriss. The research was conducted at the Centre for Integrative Neuroscience and Neurodynamics (CINN) at the University of Reading. The author wishes to thank: (1) Carien van Reekum for the use of lab facilities and feedback, (2) Francesco Saldarini, Alberto Dalla Verde and Kelsey Britton for their help in data collection and marking, and (3) Michael Lindner for help with Matlab scripts. Furthermore, the author would like to thank the participants who took part in this study. The author states no conflict of interest. To access the data, please contact Dr. Jayne Morriss.

## Footnotes

1 Surprisingly, STAI was also related to SCR magnitude during extinction [Stimulus x Time x STAI interaction: *F*(1,160.257) = 5.605, *p* =.019]. However, this effect was not significant when STAI was entered into the model alone [Stimulus x Time x STAI interaction: *F*(1,159.442) = 1.732, *p* =.190]. Similarly, when IU was entered into the model alone, the strength of the interaction dropped, however to a lesser extent [Stimulus x Time x IU interaction: *F*(1,161.292) = 2.832, *p* =.094]. Given recent replications of the effects of IU on extinction, the IU effect on SCR magnitude in this study is likely genuine, albeit underpowered.

2 Both IU and STAI were related to SCR magnitude during the picture task [Picture x IU interaction: F(1, 151.603) = 4.864, p =.029; Picture x STAI interaction: F(1, 151.603) = 7.092, p =.009]. However, both of these effects were not significant when IU and STAI were entered into the model alone [Picture x IU interaction: F(1, 141.884) = 2.160, p =.110; Picture x STAI interaction: F(1, 147.366) = 3.161, p =.077]. Thus, these effects may have been spurious.

3 The opposite pattern was observed with STAI [Levels of Uncertainty x STAI interaction: *F*(1, 193.140) = 3.381, *p* =.019]. However, this effect was not significant when STAI was entered into the model alone [Levels of Uncertainty x STAI interaction: *F*(1, 192.448) = .683, *p* =.563]. When IU was entered into the model alone the interaction remained significant [Levels of Uncertainty x IU interaction: *F*(1, 193.170) = 4.036, *p* =.008].

## References

Ben-Shakhar, G. (1985). Standardization within individuals: A simple method to neutralize individual differences in skin conductance. Psychophysiology, 22(3), 292–299.

Bennett, K. P., Dickmann, J. S., & Larson, C. L. (2018). If or when? Uncertainty’s role in anxious anticipation. Psychophysiology, e13066.

Bradley, M. M., & Lang, P. J. (2000). Emotion and motivation. Handbook of psychophysiology, 2, 602–642.

Carleton, R. N. (2016a). Fear of the unknown: One fear to rule them all? Journal of Anxiety Disorders, 41, 5–21.

Carleton, R. N. (2016b). Into the unknown: A review and synthesis of contemporary models involving uncertainty. Journal of Anxiety Disorders, 39, 30–43.

Carleton, R. N., Duranceau, S., Shulman, E. P., Zerff, M., Gonzales, J., & Mishra, S. (2016). Self-reported intolerance of uncertainty and behavioural decisions. Journal of Behavior Therapy and Experimental Psychiatry, 51, 58–65.

Chin, B., Nelson, B. D., Jackson, F., & Hajcak, G. (2016). Intolerance of uncertainty and startle potentiation in relation to different threat reinforcement rates. International Journal of Psychophysiology, 99, 79–84.

Dawson, M. E., Schell, A. M., & Filion, D. L. (2000). The Electrodermal System. In J. T. Cacioppo, L. G. Tassinary, & G. G. Berntson (Eds.), Handbook of Physiology (2nd ed., pp. 200–223). Cambridge, UK: Cambridge University Press.

Dunsmoor, J. E., Campese, V. D., Ceceli, A. O., LeDoux, J. E., & Phelps, E. A. (2015). Novelty-facilitated extinction: providing a novel outcome in place of an expected threat diminishes recovery of defensive responses. Biological Psychiatry, 78(3), 203–209.

Fergus, T. A., Bardeen, J. R., & Wu, K. D. (2013). Intolerance of uncertainty and uncertainty-related attentional biases: evidence of facilitated engagement or disengagement difficulty? Cognitive Therapy and Research, 37(4), 735–741.

Fergus, T. A., & Carleton, R. N. (2016). Intolerance of uncertainty and attentional networks: Unique associations with alerting. Journal of Anxiety Disorders, 41, 59–64.

Freeston, M. H., Rhéaume, J., Letarte, H., Dugas, M. J., & Ladouceur, R. (1994). Why do people worry? Personality and Individual Differences, 17(6), 791–802.

Fridlund, A. J., & Cacioppo, J. T. (1986). Guidelines for human electromyographic research. Psychophysiology, 23(5), 567–589.

Gentes, E. L., & Ruscio, A. M. (2011). A meta-analysis of the relation of intolerance of uncertainty to symptoms of generalized anxiety disorder, major depressive disorder, and obsessive–compulsive disorder. Clinical Psychology Review, 31(6), 923–933.

Gole, M., Schäfer, A., & Schienle, A. (2012). Event-related potentials during exposure to aversion and its anticipation: the moderating effect of intolerance of uncertainty. Neuroscience Letters, 507(2), 112–117.

Grös, D. F., Antony, M. M., Simms, L. J., & McCabe, R. E. (2007). Psychometric properties of the State-Trait Inventory for Cognitive and Somatic Anxiety (STICSA): comparison to the State-Trait Anxiety Inventory (STAI). Psychological assessment, 19(4), 369.

Grupe, D. W., & Nitschke, J. B. (2011). Uncertainty is associated with biased expectancies and heightened responses to aversion. Emotion, 11(2), 413.

Grupe, D. W., & Nitschke, J. B. (2013). Uncertainty and anticipation in anxiety: an integrated neurobiological and psychological perspective. Nature Reviews Neuroscience, 14(7), 488–501.

Hartley, C. A., & Phelps, E. A. (2012). Anxiety and decision-making. Biological Psychiatry, 72(2), 113–118.

Insel, T., Cuthbert, B., Garvey, M., Heinssen, R., Pine, D. S., Quinn, K., … Wang, P. (2010). Research domain criteria (RDoC): toward a new classification framework for research on mental disorders: Am Psychiatric Assoc.

Jacoby, R. J., Abramowitz, J. S., Buck, B. E., & Fabricant, L. E. (2014). How is the Beads Task related to intolerance of uncertainty in anxiety disorders? Journal of Anxiety Disorders, 28(6), 495–503.

Jacoby, R. J., Abramowitz, J. S., Reuman, L., & Blakey, S. M. (2016). Enhancing the ecological validity of the Beads Task as a behavioral measure of intolerance of uncertainty. Journal of Anxiety Disorders, 41, 43–49.

Jacoby, R. J., Reuman, L., Blakey, S. M., Hartsock, J., & Abramowitz, J. S. (2017). “What if I make a mistake?”: Examining uncertainty-related distress when decisions may harm oneself vs. others. Journal of Obsessive-Compulsive and Related Disorders.

Ladouceur, R., Talbot, F., & Dugas, M. J. (1997). Behavioral expressions of intolerance of uncertainty in worry: Experimental findings. Behavior Modification, 21(3), 355–371.

Lang, P. J., & Bradley, M. M. (2010). Emotion and the motivational brain. Biological Psychology, 84(3), 437–450.

Lang, P. J., Bradley, M. M., & Cuthbert, B. N. (2005). International affective picture system (IAPS): Affective ratings of pictures and instruction manual: NIMH, Center for the Study of Emotion & Attention.

Lang, P. J., Greenwald, M. K., Bradley, M. M., & Hamm, A. O. (1993). Looking at pictures: affective, facial, visceral, and behavioral reactions. Psychophysiology, 30(3), 261–273.

Lonsdorf, T. B., Menz, M. M., Andreatta, M., Fullana, M. A., Golkar, A., Haaker, J., … Kruse, O. (2017). Don’t fear ‘fear conditioning’: Methodological considerations for the design and analysis of studies on human fear acquisition, extinction, and return of fear. Neuroscience & Biobehavioral Reviews, 77, 247–285.

Lonsdorf, T. B., & Merz, C. J. (2017). More than just noise: Inter-individual differences in fear acquisition, extinction and return of fear in humans-Biological, experiential, temperamental factors, and methodological pitfalls. Neuroscience & Biobehavioral Reviews, 80, 703–728.

Lucas, K., Luck, C. C., & Lipp, O. V. (2018). Novelty-facilitated extinction and the reinstatement of conditional human fear. Behaviour Research and Therapy, 109, 68–74.

Luhmann, C. C., Ishida, K., & Hajcak, G. (2011). Intolerance of uncertainty and decisions about delayed, probabilistic rewards. Behavior Therapy, 42(3), 378– 386.

McEvoy, P. M., & Mahoney, A. E. (2012). To be sure, to be sure: Intolerance of uncertainty mediates symptoms of various anxiety disorders and depression. Behavior Therapy, 43(3), 533–545.

Morriss, J., Chapman, C., Tomlinson, S., & Van Reekum, C. M. (2018). Escape the bear and fall to the lion: The impact of avoidance availability on threat acquisition and extinction. Biological Psychology, 138, 73–80.

Morriss, J., Christakou, A., & Van Reekum, C. M. (2015). Intolerance of uncertainty predicts fear extinction in amygdala-ventromedial prefrontal cortical circuitry. Biology of Mood & Anxiety Disorders, 5(1), 1.

Morriss, J., Christakou, A., & Van Reekum, C. M. (2016). Nothing is safe: Intolerance of uncertainty is associated with compromised fear extinction learning. Biological Psychology, 121, 187–193.

Morriss, J., Gell, M., & van Reekum, C. M. (2018). The uncertain brain: A co-ordinate based meta-analysis of the neural signatures supporting uncertainty during different contexts. Neuroscience & Biobehavioral Reviews, 96, 241–249.

Morriss, J., Macdonald, B., & van Reekum, C. M. (2016). What Is Going On Around Here? Intolerance of Uncertainty Predicts Threat Generalization. PloS one, 11(5), e0154494.

Schienle, A., Köchel, A., Ebner, F., Reishofer, G., & Schäfer, A. (2010). Neural correlates of intolerance of uncertainty. Neuroscience letters, 479(3), 272–276.

Shihata, S., McEvoy, P. M., Mullan, B. A., & Carleton, R. N. (2016). Intolerance of uncertainty in emotional disorders: What uncertainties remain? Journal of Anxiety Disorders, 41, 115–124.

Solnik, S., DeVita, P., Rider, P., Long, B., & Hortobágyi, T. (2008). Teager–Kaiser Operator improves the accuracy of EMG onset detection independent of signal-to-noise ratio. Acta of bioengineering and biomechanics/Wroclaw University of Technology, 10(2), 65.

Somerville, L. H., Wagner, D. D., Wig, G. S., Moran, J. M., Whalen, P. J., & Kelley, W. M. (2013). Interactions between transient and sustained neural signals support the generation and regulation of anxious emotion. Cerebral Cortex, 23(1), 49–60.

Spielberger, C. D., Gorsuch, R. L., Lushene, R., Vagg, P., & Jacobs, G. (1983). Consulting Psychologists Press, Inc. 2». Palo Alto (CA).

Tanovic, E., Gee, D. G., & Joormann, J. (2018). Intolerance of uncertainty: Neural and psychophysiological correlates of the perception of uncertainty as threatening. Clinical Psychology Review.

Tanovic, E., Hajcak, G., & Joormann, J. (2018). Hating waiting: Individual differences in willingness to wait in uncertainty. Journal of Experimental Psychopathology, 9(1), 2043808718778982.

Tanovic, E., Pruessner, L., & Joormann, J. (2018). Attention and anticipation in response to varying levels of uncertain threat: An ERP study. Cognitive, Affective, & Behavioral Neuroscience, 1–14.

Tassinary, L. G., Cacioppo, J. T., & Vanman, E. J. (2007). The skeletomotor system: Surface electromyography. In J. T. Cacioppo, L. G. Tassinary, & G. G. Berntson (Eds.), Handbook of psychophysiology (pp. 267–299). New York, NY, US: Cambridge University Press.

